# Ultrasound-mediated mechanical forces selectively kill tumor cells

**DOI:** 10.1101/2020.10.09.332726

**Authors:** Ajay Tijore, Felix Margadant, Mingxi Yao, Anushya Hariharan, Claire Alexandra Zhen Chew, Simon Powell, Glenn Kunnath Bonney, Michael Sheetz

## Abstract

Ultrasound has been used to target tumors either through local heating or local nanobubbles but these methods damage surrounding normal cells in the target area. Recent studies show that tumor cells are susceptible to mechanical stresses and undergo calcium-dependent apoptosis under conditions that promote normal cell growth. Here we report that low-frequency ultrasound causes apoptosis of tumor cells by activating a calpain-dependent mitochondrial pathway that depends upon calcium entry through the mechanosensitive Piezo1 channels. This is a general property of all tumor cell lines tested so far irrespective of tissue origin. In animals, ultrasound irradiation causes tumor killing in the chick chorioallantoic membrane (CAM) model with relatively little damage to the chick embryos. Further, patient-derived pancreatic tumor organoids are killed by ultrasound treatment. Because low-level ultrasound causes apoptosis of tumor cells from many different tissues in different microenvironments, it may offer a safe non-invasive approach to augment tumor treatments.

## Introduction

Mechanical perturbations of tumor cells appear to increase apoptosis or inhibit tumor growth in multiple recent studies. For example, physiologically relevant fluid shear forces caused apoptosis in circulating as well as adherent tumor cells.^1, 2^ In a mouse model, exercise or stretching of tumors inhibits tumor growth.^3, 4^ Further, mechanically active muscle tissue has a low risk of tumor formation. In fact, muscle-associated tumors doesn’t even make it to the list of the most commonly occurring 36 tumors worldwide.^5^ With regard to mechanism, our earlier studies document the mechanism of tumor cell growth inhibition and apoptosis by mechanical stress.^6^ The stretch-mediated apoptosis depends on the mechanosensitive Piezo1 channel activity that enables calcium entry upon mechanical activation, initiating a process of calpain-dependent apoptosis. In a broader context, normal cells can become transformed like tumor cells upon the depletion of rigidity sensing activity by diminution of protein components of the rigidity sensor.^7, 8^ Surprisingly, such transformed normal cells from several different tissues become mechanosensitive and apoptose after stretching.^6^ Since the majority if not all tumor cells are depleted of rigidity sensors, mechanosensitivity appears to be a general feature of the tumor cells irrespective of tissue origin.^9^

In the past, ultrasound has been used to sensitize tumor cells in combination with different strategies such as sonodynamic therapy^10^, chemotherapy^11^ and hyperthermia^12^. In addition, different ultrasound modes have been tested individually to treat tumors using high-intensity focused ultrasound (HIFU)^13^, high-intensity pulsed ultrasound^14^ and low-intensity pulsed ultrasound^15^. However, concerns were raised about healthy tissue damage surrounding the target area and hence these methods found limited clinical use. Since many tumor cells appear mechanosensitive, we decided to test the effect of ultrasound-mediated mechanical forces that do not heat or otherwise damage healthy cells. In this study, we use low-frequency ultrasound under conditions that do not damage the normal cells but penetrate tissue and apply mechanical stresses on cells through pressure pulses.^16^ Many applications for low-frequency ultrasound have been approved for human use and there is little evidence of damage to normal tissue by ultrasound.^17–19^ Consistent with those observations, we find that tumor cells, tumors and tumor organoids are killed by ultrasound in a frequency range and at power levels that are approved for human use.^19, 20^

## Results

### 1. Ultrasound induces tumor cell apoptosis

First, we tested the efficacy of low-frequency ultrasound to induce tumor cell apoptosis *in vitro*. Tumor cells were grown on matrigel and subjected to low-frequency ultrasound (33 kHz) for 2 hr with ~7.5 W power level (total output). A significant increase in apoptosis (46% for MDA-MB-231, 58% for A375p and 32% HT1080) was observed with 50% duty cycle in comparison to the non-treated tumor cells (9% for MDA-MB-231, 4% for A375p and 5% HT1080) (Figure 1a & b). In contrast, when normal cells from the same tissue of origin as the tumor cells were subjected to ultrasound treatment, negligible apoptosis was observed (16% for MCF10A and 3% for HFF after ultrasonication *vs.* 15% for MCF10A and 4% for HFF for non-treated controls).

**Figure 1.**
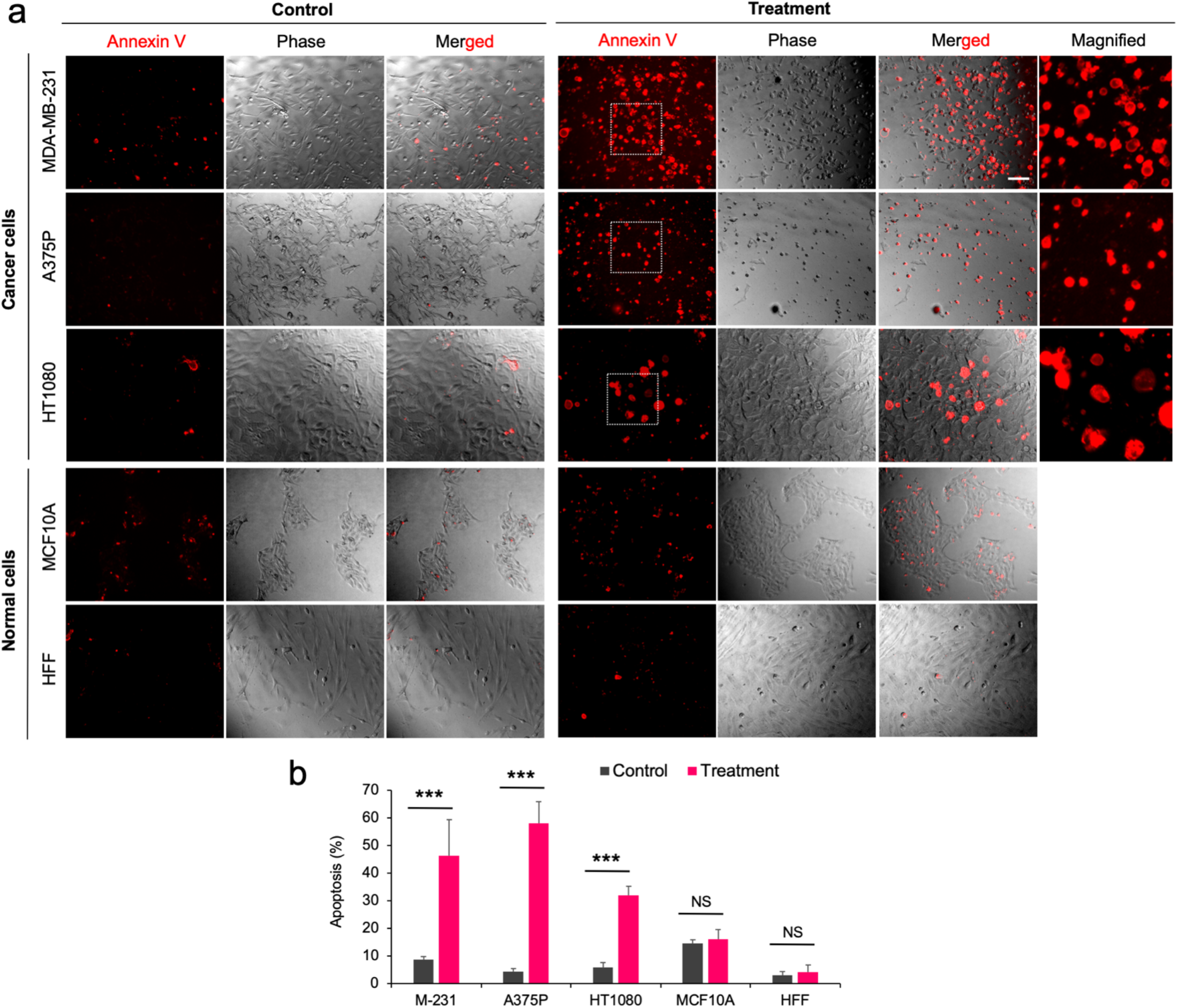
Ultrasound treatment promotes apoptosis in tumor cells grown on matrigel. **a)** Image panels showing apoptosis level in tumor cells (MDA-MB-231, A375p, HT1080) and normal cells (MCF10A, HFF) with and without ultrasound treatment, scale bar: 50 μm. Magnified view of apoptotic cells were shown separately. **b)** Bar diagram illustrating apoptosis level in tumor and normal cells with and without ultrasound treatment. n>600 cells, data are representative of two independent experiments, Annova test, *p*\*\*\*< 0.001.

Next, we assessed the ability of ultrasound to inflict the necrosis. About 8-9% tumor cells (MDA-MB-231 and A375p) displayed both apoptosis plus necrosis upon treatment in comparison to 32-35% cells showing only apoptosis (Figure S1). To understand the role of ultrasound frequency in promoting apoptosis, experiments were repeated using another ultrasound frequency (120 kHz) with the same power level settings. However, there was no increase in tumor cell apoptosis upon treatment unlike with 33 kHz frequency (Figure S2). To test the generality of killing, tumor cells from another tissue (SKOV3 ovarian adenocarcinoma) were treated with ultrasound. An increase in apoptosis (32%) was seen in comparison to the non-treated tumor cells (7%) (Figure S3). Thus, results showed that low-frequency ultrasound caused tumor cell apoptosis compared to relatively high-frequency ultrasound in many tumor cell lines, which was reminiscent of the effects of cyclic mechanical stretch.^6^

### 2. Repetitive ultrasound exposure results in similar level of apoptosis each time

To test if repeated ultrasound treatments would produce an ultrasound-resistant population of tumor cells, cells were treated with ultrasound for 2 hr/day for three consecutive days. Viable cells were followed with calcein AM dye. A marked decrease in the tumor cell population (MDA-MB-231 or A375p) was observed after each round of treatment with no indication of a plateau. Further, the number of remaining viable cells decreased in an apparently exponential manner (Figure 2a). In contrast, non-treated tumor cells showed a progressive growth with time. Treatment of normal cells (MCF10A and HFF) caused no significant change in growth rate compared to non-treated cells (Figure 2a). Further, fluorescence intensity measurements showed a significant decrease in tumor cell population after treatment in comparison to the non-treated tumor cells and both normal cell groups (Figure 2b). Thus, with successive ultrasound treatment, there was a similar level of tumor cell apoptosis each time, producing a dramatic overall decrease in tumor cell number irrespective of tumor origin.

**Figure 2.**
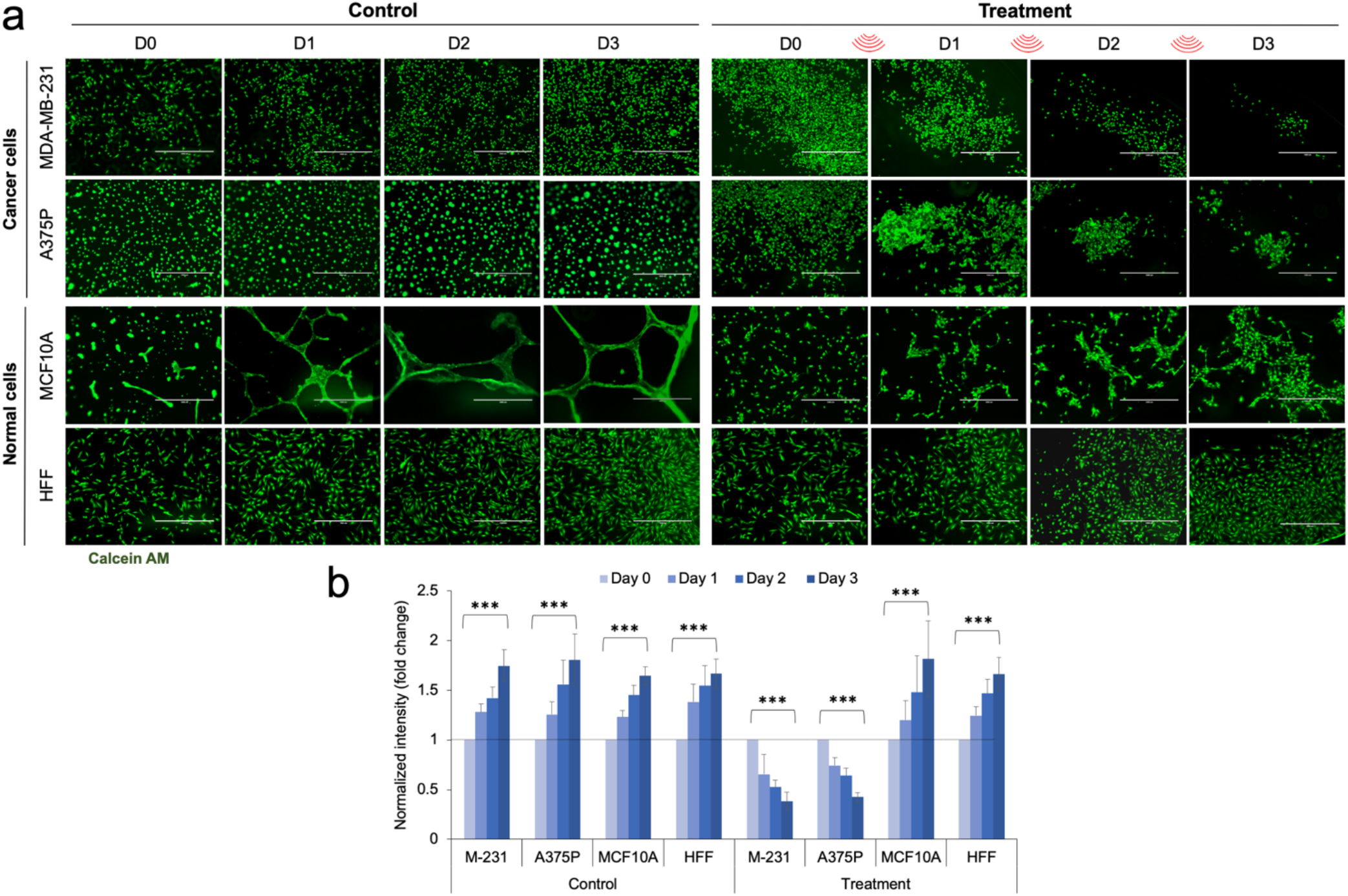
Repetitive ultrasound exposure reduces tumor cell growth on matrigel. **a)** Panels showing viable tumor and normal cells with and without repetitive ultrasound treatment from day 0 to day 3. Calcein AM was used to stain viable cells. Scale bar: 1000 μm. **b)** Bar diagram showing normalized fluorescence intensity in tumor and normal cells in the control and ultrasound-treated cells from day 0 to day 3. Day 0 intensity was considered one and shown by dotted line. n>5 image fields, data are representative of two independent experiments, Annova test, *p*\*\*\*< 0.001.

### 3. Piezo1 channel mediated-calcium influx is required for tumor cell apoptosis

The stretch-induced tumor cell apoptosis (mechanoptosis) relied upon the mechanosensitive Piezo1 channels that catalyzed calcium entry^21^ and we postulated that ultrasound-induced apoptosis involved a similar mechanism.^6^ To test the role of Piezo1 in ultrasound-mediated apoptosis, Piezo1 knock-down (KD) cells were treated with ultrasound and the apoptosis level was measured (Figure 3a and 3b). Interestingly, the apoptosis level was low in Piezo1 KD cells compared to control siRNA cells upon ultrasound treatment (Figure 3c). Thus, as found for stretch-induced apoptosis, Piezo1 was also required for the ultrasound-based mechanoptosis.

**Figure 3.**
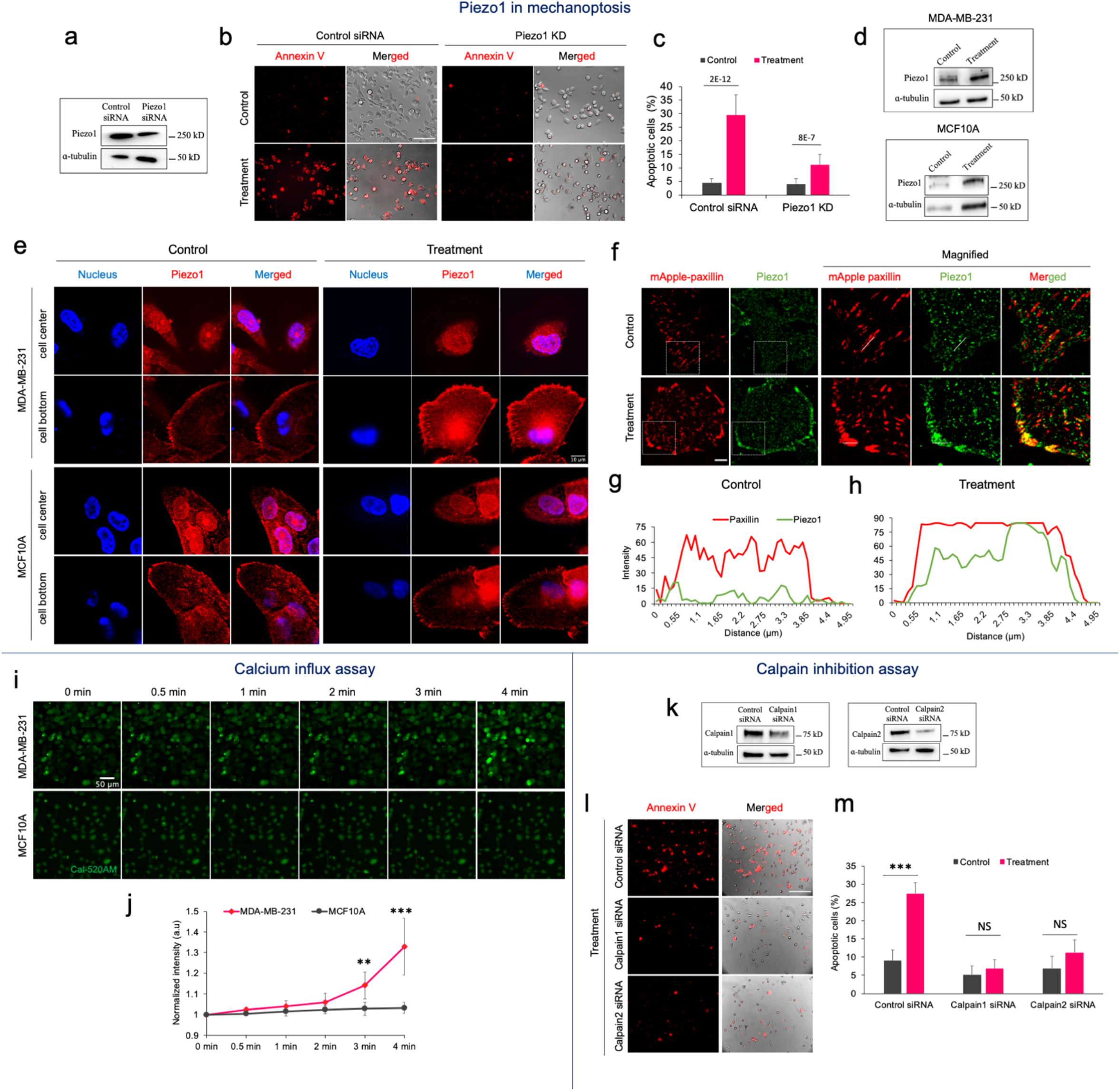
Ultrasound enhances calcium influx through activated Piezo1 channels in tumor cells. **(a)** Western blot showing Piezo1 expression in control siRNA-and Piezo1 siRNA-treated MDA-MB-231 cells. **(b)** Representative images displaying apoptotic cells with and without US treatment in control siRNA- and Piezo1 siRNA-treated tumor cells. Scale bar: 100 μm. **(c)** Bar diagram exhibiting level of apoptosis in tumor cells under different conditions, n>1000 cells, student t-test, *p*\*\*\*<0.001. **(d)** Western blots showing Piezo1 expression in MDA-MB-231 and MCF10A cells with and without US treatment. **(e)** Confocal images displaying spatial distribution of Piezo1 at the cell centre and bottom in MDA-MB-231 and MCF10A cells with and without US treatment. Scale bar: 10 μm. **(f)** Representative images showing spatial distribution of paxillin and Piezo1 in MDA-MB-231 cells with and without ultrasound treatment. Scale bar: 10 μm. **(g-h)** Fluorescent intensity plot displaying paxillin and Piezo1 distribution in cells with and without ultrasound treatment. **(i)** The time-lapse montage displaying Cal-520 AM dye intensity upon treatment in MDA-MB-231 and MCF10A cells. Scale bar 50 μm. **(j)** Graph representing Cal520 AM dye intensity plot of tumor and normal cells, n= 5 image fields from two experiments, student t-test, *p*\*\*<0.01, *p*\*\*\*<0.001. **(k)** Western blot showing calpain1 and 2 expression after siRNA knockdown in tumor cells. **(l)** Representative images showing apoptotic tumor cells with ultrasound treatment in the presence of control siRNA, calpain1 siRNA and calpain2 siRNA. Scale bar: 100 μm. **(m)** Bar diagram displaying the extent of apoptosis in presence of different siRNAs with and without ultrasound treatment. n>1000 cells, data are representative of two experiments, student t-test, *p*\*\*\*<0.001.

Although there was a similar expression level of Piezo1 in different tumor cells and their matched normal counterparts^6^ (Figure S4), there was a possibility that Piezo1 level increased with ultrasound treatment. To test this possibility, cells (MDA-MB-231 and MCF10A) were treated with ultrasound for 2 hr. Surprisingly, western blot results showed an increase in Piezo1 expression in tumor cells upon ultrasound treatment but not in normal cells (Figure 3d). Further, to elucidate ultrasound-mediated precise changes in spatial distribution of Piezo1, high-resolution imaging was performed. Interestingly, distinct Piezo1 localization to the plasma membrane was observed in tumor cells following ultrasound treatment, whereas it remained diffuse in the cytoplasm of non-treated tumor cells (Figure 3e). In stark contrast, in normal cells, Piezo1 was predominately located in the nuclear region (especially nuclear envelop)^22^ with and without ultrasound treatment. In particular, no redistribution of Piezo1 to the plasma membrane was detected in normal cells upon treatment unlike tumor cells. These observations were further confirmed by wide-field imaging. In particular, a 55% reduction in the nuclear to cytoplasmic intensity ratio was observed in treated tumor cells compared to non-treated ones, while in normal cells, there was 29% reduction in the ratio upon treatment (Figure S5). Recent studies have reported the association of Piezo1 with peripheral mature adhesion.^23, 24^ Hence, we examined if ultrasound treatment promoted Piezo1 translocation to the peripheral adhesions located in the plasma membrane of tumor cells. Cells transiently transfected with mApple-paxillin were used to determine the adhesion distribution. In control tumor cells, Piezo1 was mostly diffuse in the cytoplasm with no sign of association with peripheral adhesions (Figure 3f & 3g). In contrast, ultrasound treated tumor cells showed strong association of Piezo1 with the enlarged adhesions at cell periphery (Figure 3f & 3h). Thus, results demonstrated that ultrasound treatment promoted Piezo1 expression as well as its localization to the mature adhesions at cell periphery in tumor cells, while it was primarily confined to the nuclear region in normal cells.

Next, based on above results, we postulated that ultrasound may trigger calcium influx through Piezo1 localized at plasma membrane in tumor cells. In contrast, the nuclear distribution of Piezo1 in normal cells was consistent with the possibility that Piezo1 may not alter calcium entry through plasma membrane upon ultrasound treatment. To test our hypothesis, an ultrasound-induced calcium influx assay was performed. To quantify the intracellular calcium level, cells were loaded with calcium sensitive dye Cal-520AM and subjected to the ultrasound treatment (Figure 3i). In tumor cells, a significant increase in the fluorescence intensity was observed 3 min after the treatment; but no such increase in the intensity was observed in normal MCF10A cells (Figure 3j). The role of calpain in mitochondria-dependent apoptosis has been widely established.^6, 25^ Hence we decided to test the role of calcium activated-calpain protease in mechanoptosis. To do that, siRNAs against calpain 1 and 2 were used to prepare calpain KD tumor cells (Figure 3k). Cells treated with scrambled siRNA were used as the control. After ultrasound treatment, both calpain 1 and 2 knockdown cells had remarkably low levels of apoptosis compared to the ultrasound-treated control cells (Figure 3l and 3m), indicating that calpain played major role in mechanoptosis. Collectively, the results demonstrated that ultrasound initiated-calcium influx through activated Piezo1 channels acted as an upstream inducer of mechanoptosis. Calcium activated-calpain subsequently triggered mitochondria-mediated apoptosis through a chain of reactions as described previously.^6^ In stark contrast, normal cells were unaffected by ultrasound-mediated mechanical forces because ultrasound did not cause a dramatic rise in cytosolic calcium level.

### 4. Ultrasound impedes *in vivo* tumor growth in the CAM tumor model

Since ultrasound killed tumor cells grown on the matrigel, we hypothesized that similar ultrasound treatment would promote tumor killing *in viv*o without causing noticeable damage to healthy tissues. To test the hypothesis, we inoculated chick chorioallantoic membranes (CAM) with stable GFP-transfected MDA-MB-231 tumor cells in a matrigel plug and allowed them to grow for two days. After tumors were established (often with vascularization), the embryos were ultrasonicated for 2 hr/day for two consecutive days (Figure 4a and Figure S6). Consistent with the *in vitro* results, we found an average two-fold reduction in GFP fluorescence intensity after two days, indicating cell killing in the treated tumors. In contrast, non-treated tumors displayed an increase in the intensity, indicating normal tumor growth (Figure 4b and 4c). Next, we performed annexin V-based apoptosis assay on carefully excised ultrasound-treated and control tumors. Consistent with *in vitro* results, treated tumors had a high fraction of apoptotic cells (Figure 4d and 4e). However, in the control tumors, few apoptotic cells were observed. In addition, the viability of chick embryos was assessed after every round of ultrasound treatment. There was a 100% survival rate after the first round of ultrasound (Figure 4f). After the second round of ultrasound, chick embryo survivability was reduced to 60% while, it remained 100% in untreated embryos. Thus, these findings indicated that ultrasound treatment killed tumor cells in the chick embryo while, causing less damage to delicate embryonic tissues/organs.

**Figure 4.**
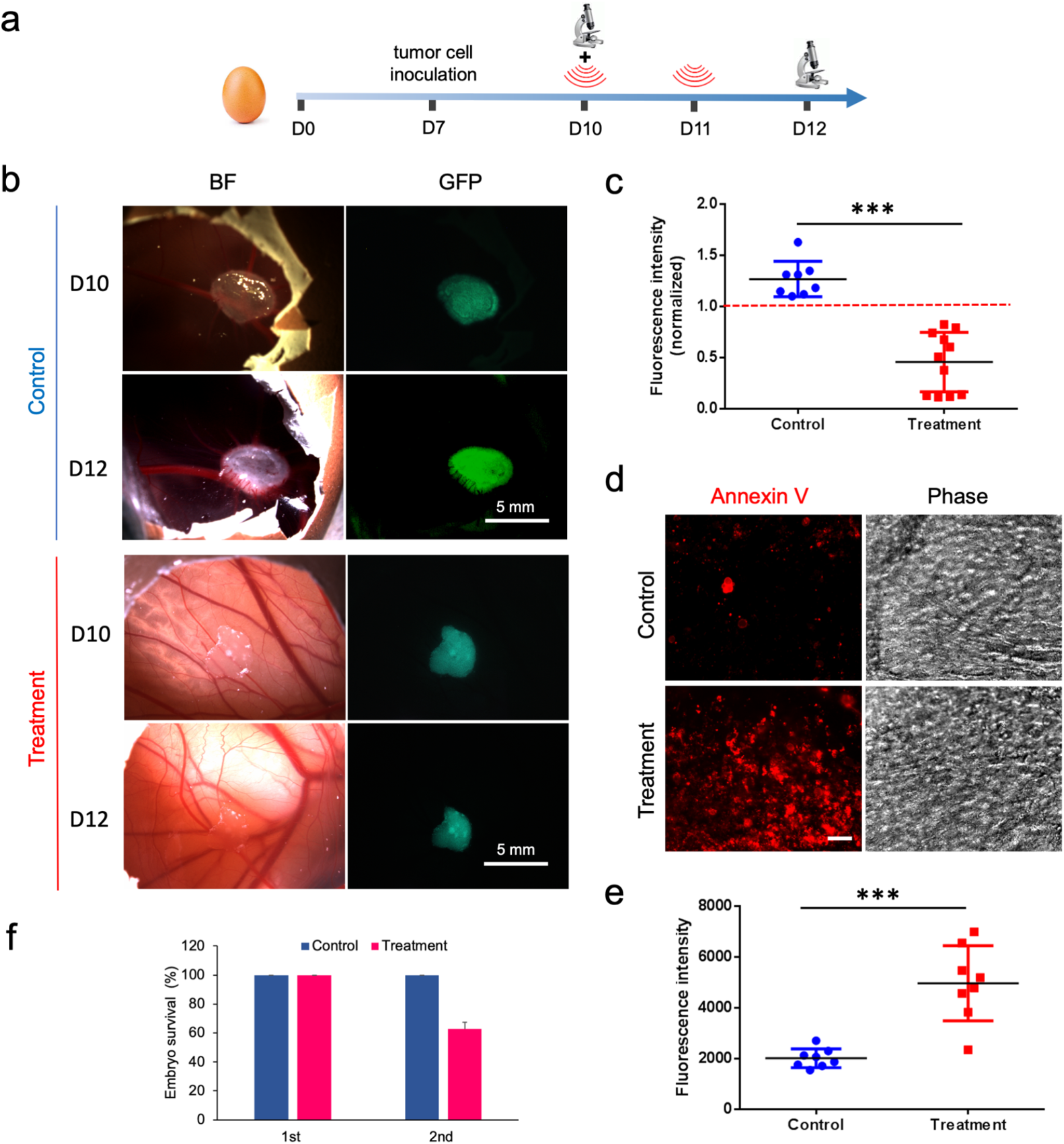
Ultrasound causes reduction in MDA-MB-231 tumor growth in CAM model. **a)** Time-line depicting the scheme of ultrasonicating the tumors grown in chick embryo model. **b)** Top two panel rows display bright-field and GFP expression images of tumor before (D10) and after two rounds (D12) of ultrasound treatment. Bottom panel rows show bright-field and GFP expression image of tumor without any treatment on D10 and D12. **c)** Graph showing fluorescence intensity of tumors with and without ultrasound treatment on D12. Baseline intensity is shown by red dotted line. (n=8 for control and n= 11 for treatment), data are representative of three independent experiments, student t-test, *p*\*\*\*< 0.001. **d)** Image panels showing annexin-V stained apoptotic cells in CAM tumors with and without treatment. **e)** Graph showing annexin-V intensity of CAM tumors with and without ultrasound treatment. (n=8 image fields from control and treated samples), student t-test, *p*\*\*\*< 0.001. **f)** Bar diagram showing chick embryo survival rate after two rounds of ultrasound treatment (n=13).

To determine if other transformed cells behaved similarly, we used HEK293T cells that were stably transformed by SV40 large T antigen. We selected HEK293T cells because these were p53 mutated cells that exhibit transformed growth on soft agar, showed tumorigenic potential (form tumors in nude mice) and had high transfection efficacy.^26^ Consistent with previous results, there was a significant reduction in GFP fluorescence intensity in ultrasound-treated tumors whereas, non-treated tumors showed an increase in the intensity (Figure S7). Thus, the ultrasound mechanosensitivity was not limited to a specific tumor or tumor cell line.

### 5. Ultrasound causes apoptosis of patient-derived tumor organoids

To further validate the ultrasound-induced mechanoptosis, human pancreatic tumor organoids were subjected to the ultrasound treatment. Organoids were derived from primary tissue/tumor biopsies and were composed of organ-specific cell type aggregates that were capable of self-renewal, self-organization and retained organ functionality.^26, 27^ Freshly prepared organoids were treated with ultrasound for 2 hr/day for two consecutive days (Figure 5a). On Day 3, there was a marked increase in the apoptosis level in treated tumor organoids in comparison to the controls (Figure 5b and 5c). In addition, many organoids were disrupted and had a broken and irregular morphology in the treated samples (Figure 5d and Figure S8). These disrupted organoids were surrounded by many isolated cells/aggregates which exhibited high level of apoptosis. The remaining organoids in the treated sample were significantly smaller (~7500 um²) when compared with the controls (~44000 um²) (Figure 5f and Figure S8). It seems that disrupted organoids reorganized to form smaller organoids. Next, F-actin imaging revealed no visible difference in stress fibre assembly in the treated and control groups (Figure 5e). Thus, the results illustrated that the human-derived tumor cells in a organoid that approximated the organization of *in vivo* tumor were vulnerable to the ultrasound-mediated mechanical forces like the established tumor cell lines.

**Figure 5.**
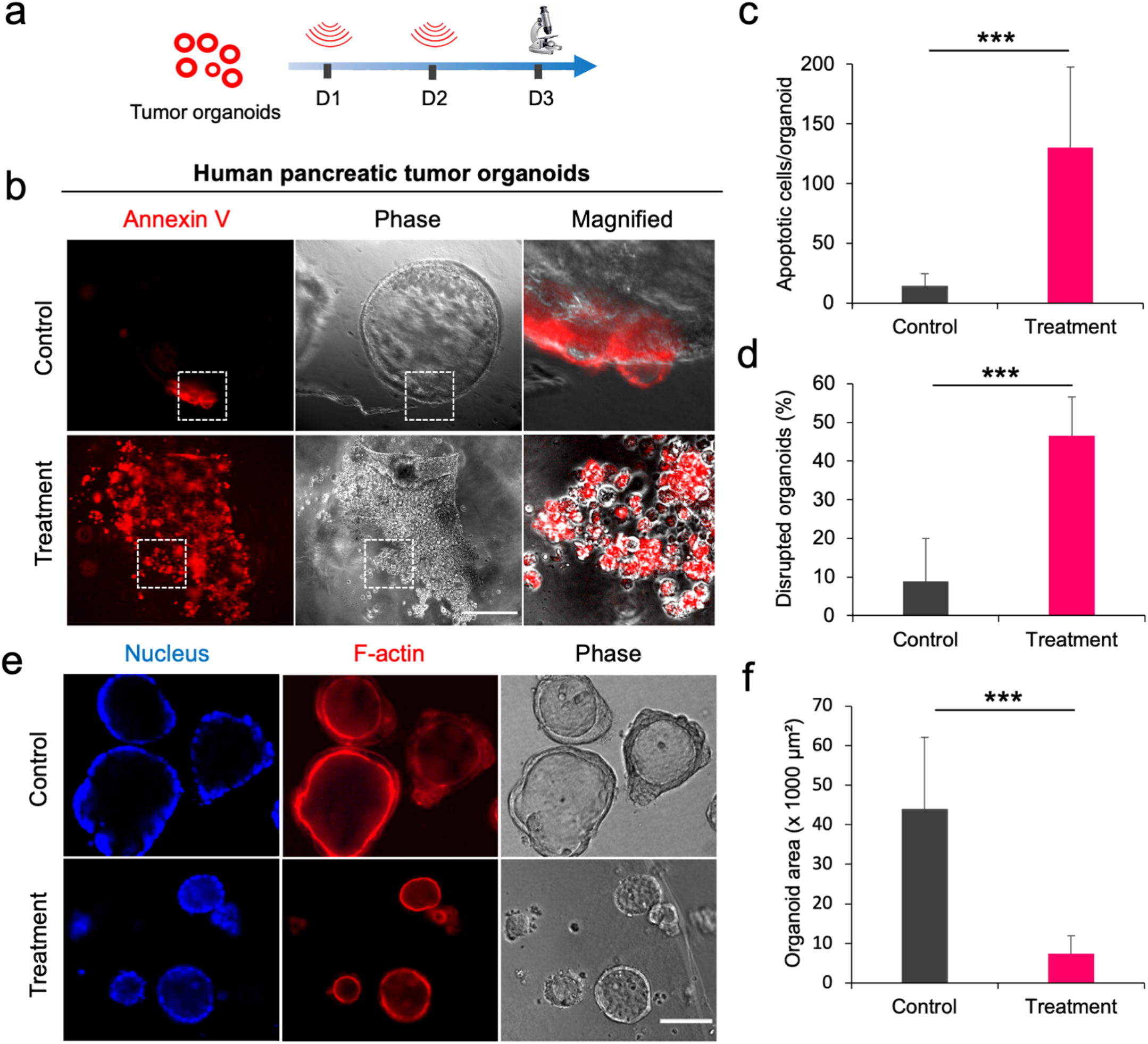
Ultrasound promotes apoptosis in human pancreatic tumor organoids. **a)** Timeline depicting the scheme of tumor organoid US treatment. **b)** Representative images showing apoptotic cells/organoids in control and treated samples after two rounds of ultrasound, scale bar: 100 μm. **c)** Bar diagram showing the extent of apoptosis per organoids in the control and treated organoids. n>25 organoids, student t-test, *p*\*\*\*< 0.001. **d)** Bar diagram estimating population of disrupted organoids in the control and treated organoids. n>100 organoids, student t-test, *p*\*\*\*< 0.001. **e)** Immunofluorescence staining of F-actin and nucleus of organoids in the control and treated samples, scale bar: 100 μm. **f)** Bar diagram showing the organoid area in the control and treated organoids. n>100 organoids, student t-test, *p*\*\*\*< 0.001.

## Discussion

In these studies, we find that tumor cells are killed by low-frequency ultrasound under conditions that do not damage normal cells/tissues. The tissue of origin of the cells does not significantly affect the level of apoptosis caused by ultrasound treatment. Apoptosis requires the presence of Piezo1 at the cell periphery and appears to be mediated by mechanically-dependent calcium entry that activates calpain to cause an apoptotic cascade. The ultrasound effects on the chick CAM tumor and patient-derived tumor organoids are dramatic with significant killing of tumor cells in physiologically relevant tissue forms. Thus, we suggest that ultrasound irradiation acts at a subcellular level that is relatively insensitive to the cell microenvironment but is dependent upon the transformed state of the tumor cells.^9^

This selective killing correlates with differences in the mechanical properties of tumor and normal cells.^18^ For example, all tumor cell lines that we have tested lack rigidity sensors (e.g. tropomyosin 2.1 and myosin IIA) enabling them to grow on soft surfaces while making them sensitive to apoptosis upon cyclic stretching.^6, 8, 9^ In contrast, normal cells with rigidity sensors apoptose on soft surfaces but are not killed by cyclic stretching.^7^ Interestingly, restoration of rigidity sensors in tumor cells promotes rigidity-dependent growth, improves cell stiffness and cells even grow when subjected to cyclic stretching.^6, 8^ In contrast, depletion of rigidity sensors in normal cells initiates transformed growth as well as apoptosis upon cyclic stretching. Since we have tested cells from different tissues, these findings indicate that the great majority of tumor cells from many tissues are vulnerable to mechanical forces irrespective of their microenvironment, whereas normal cells withstand mechanical treatments and continue to grow in a proper environment.^2, 3, 28^

In the past, several attempts have been made to cause tumor cell killing with ultrasound. However, those were based on either micro/nanobubble-generated thermal cavitation, thermal ablation or focused high energy waves which are often associated with a high risk of local healthy tissue damage due to overheating.^20^ For instance, high intensity focused ultrasound (HIFU) has been used clinically to ablate tumors located within body but it’s application is limited due to severe thermal side-effects and limited ability to treat metastatic tumors.^29, 30^ In stark contrast, ultrasound treatments used in these studies caused no heating of any of the cell samples during *in vitro* or *in vivo* conditions. Hence, normal cells (MCF10A and HFF) remain unharmed upon optimized ultrasound irradiation (Figure 1 and 2). More importantly, in the chick CAM model tumor cells were killed with a lower risk to the chick embryo that is highly vulnerable to ultrasound treatment (Figure 5f).^31^ Thus, low-frequency ultrasound preferentially kills tumor cells. However, more studies are required to further optimize the selectively of the tumor cell killing.

At a mechanistic level, ultrasound-induced mechanoptosis appears to act similarly to the mechanical stretch-induced mechanoptosis.^6^ The ultrasound pressure waves appear to activate Piezo1 channels that allows calcium entry, which activates calpain and subsequently initiates a mitochondrial apoptotic pathway (Figure 6). This calcium-induced apoptosis is similar to calcium-dependent apoptosis of tumor cells through the ER-mitochondrial stress pathway that is activated by several natural compounds.^32, 33^ With a more detailed understanding of the mechanism of tumor cell death, there are many possible cancer therapies that would enhance the effect of ultrasound treatment. Because, ultrasound has been approved for human exposure at power levels ten to hundred-fold higher than the levels used in this study^20^, we suggest that it is practical to develop ultrasound-based therapies that could augment tumor treatments.

**Figure 6.**
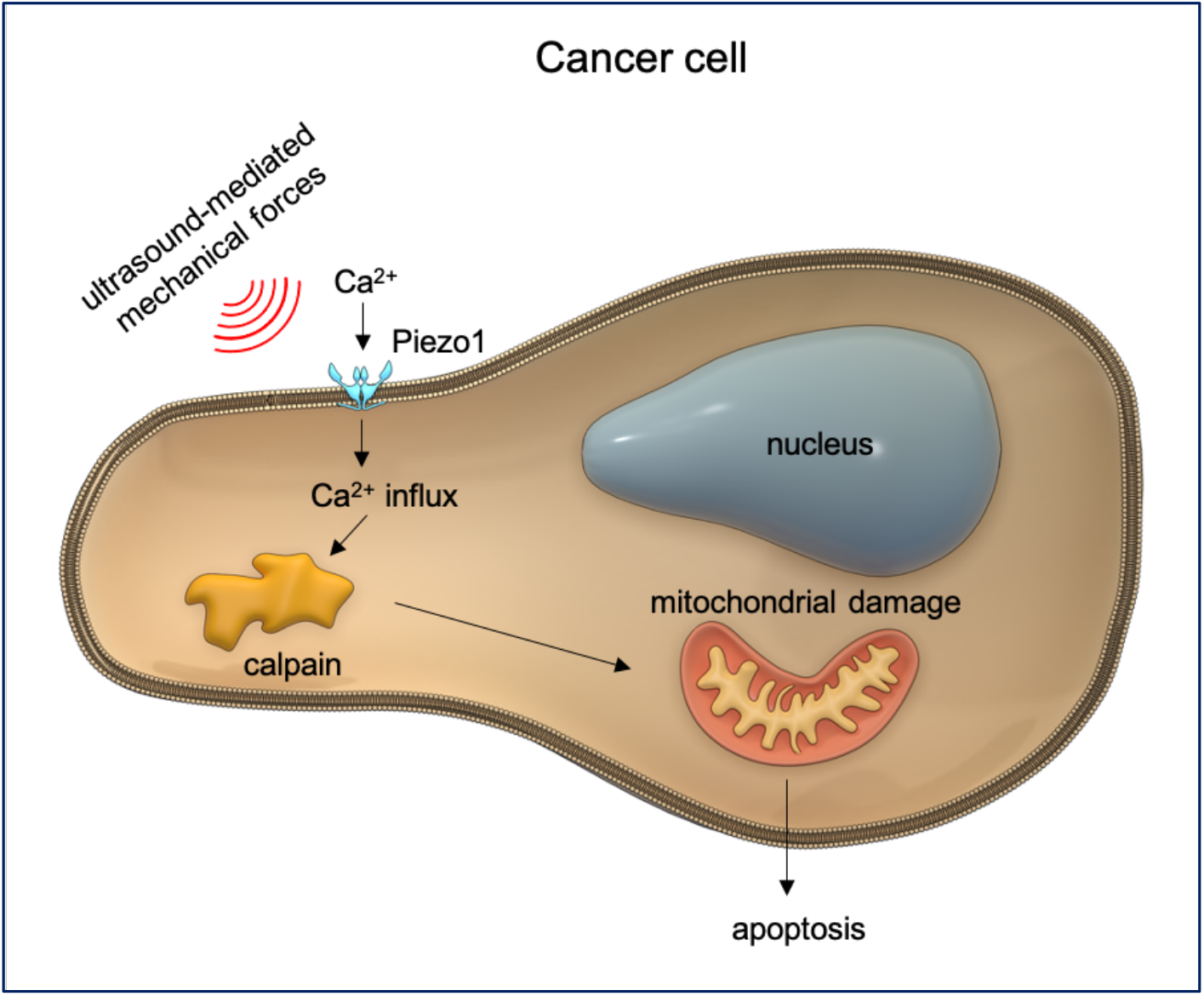
Schematic illustrating the mechanism of ultrasound-mediated tumor cell mechanoptosis. Ultrasound generated-mechanical forces activate mechanosensitive Piezo1 channels to initiate rapid calcium entry. Calcium-activated calpain then triggers mitochondrial apoptotic pathway through chain of reactions to ultimately induce mechanoptosis.

## Acknowledgement

We would like to thank all group members of Sheetz lab. A.T and M. Y are supported by the Singapore Ministry of Education Academic Research Fund Tier 3 (MOE grant No: 2016 T3-1-002). M.S is supported by NIH grants, NUS grants and Mechanobiology Institute, National University of Singapore.

## Contribution

A.T., F.M. and M.S. conceived and designed the project.; A.T. performed cell and organoid experiments; F.M. fabricated ultrasound devices; A.T. and A.H. performed CAM experiments; A.T., M.Y. and S.P. performed calcium related experiments; C.A.Z.C. and G.B. developed tumor organoids, A.T., F.M. and M.S. wrote manuscript. All authors read and commented on manuscript.

## Methods

### Cell culture

Cell lines MDA-MB-231 (gift from Dr. Jay Groves, UC Berkeley), A375p (gift from Dr. Chris Bakal, ICR UK) and primary human foreskin fibroblasts (HFF, ATCC), HT1080 (ATCC) and SKOV3 (gift from Dr. Ruby Huang, CSI, NUS) were cultured in high glucose DMEM (ThermoFisher) supplemented with 10% FBS and 1% penicillin and streptomycin at 37°C in 5% CO2 environment. MCF10A, human breast epithelial cells (ATCC) were grown in a specialized culture medium as mentioned previously.^7^ For cell experiments, high-glucose DMEM containing 10% FBS and 1% penicillin and streptomycin was utilized. For cell trypsinization, TrypLE (ThermoFisher) was used and cultured on matrigel (Corning) coated surfaces.

### Surface coating with matrigel

96 well plate surface was coated with thin layer of matrigel (Corning) according to manufacturer’s protocol. Briefly, after thawing matrigel at 4 °C, 33 μL of matrigel solution was quickly added on to pre-chilled surface of a single well of 96 well plate. Well plates were incubated at 37°C for 24-48 hr to solidify matrigel before performing a cell seeding.

### Plasmids and transfection

GFP tagged-stable MDA-MB-231 cell line was gift from Pan Meng (MBI, NUS) and used for *in vivo* study. For transient transfection of GFP and mApple-paxillin plasmid, Lipofectamine 3000 reagent (ThermoFisher) was used according to the manufacturer’s protocol.

### siRNA assay

To perform assays, cells were plated in a six well plate. Calpain1, calpain2, Piezo1 and scramble siRNA (Sigma) transfections were done on the next day using lipofectamine RNAiMAX (Invitrogen) according to the manufacturer’s protocol. Calpain siRNAs were prepared by Protein Cloning Expression Facility, MBI, NUS as described previously.^6^

### Western blot study

The cells were incubated with siRNA for 48 h, pelleted, washed with PBS. RIPA buffer (Sigma) supplemented with 1X cOmplete protease inhibitor (Roche, No. 4693116001) was used for cell lysis purpose. Cell mixture was centrifuged at 15000 rpm, 4°C for 20 min. The supernatant was transferred to eppendorf and mixed with 2X loading dye (2X Lammeli Buffer, Bio-Rad, No. 1610737) + beta-Mercaptoethanol (Sigma). Samples were denatured for 5 min at 95°C and gel was run using a 4-20% Mini-PROTEAN ® TGX precast protein gels (Bio-Rad, No. 4561094). Next, gels were then transferred onto blot paper and blocked with 5% BSA solution in 1X TBST (Tris-Buffered Saline with Tween-20) for 1 hr. The membrane was then incubated with primary antibody overnight at 4°C on shaker. Membranes were washed with TBST three times (10 min per wash) followed by incubation with secondary antibodies in 1X TBST (Horse Radish Peroxidase – HRP) for 1 hr. The chemiluminescence of the membranes was developed using Super Signal Femto Substrate Kit (Pierce) and developed using ChemiDoc Touch Imager (Bio-Rad). Primary antibodies and conditions used are as follows: Calpain 1 (rabbit, 1:1000, Abcam, ab39170), Calpain 2 (rabbit, 1:1000, Abcam, ab39165), Piezo1 (rabbit, 1:500, Novus Biologicals, NBP1-78446), alpha-tubulin (mouse, 1:3000, Sigma T9026). The secondary antibodies, HRP-conjugated goat anti-rabbit IgG (1:2000, Bio-Rad, 170-6515) and goat anti-mouse IgG (1:2000, Bio-Rad, 170-6516).

### Apoptosis and necrosis assay

To identify cell apoptosis, Annexin V-Alexa Fluor 488 or Annexin V-Alexa Fluor 594 conjugates (Thermofisher Scientific) were used according to the manufacturer’s protocol. To check cell necrosis, propidium iodide from live/dead cell double staining kit (Sigma Aldrich) was used according to the manufacturer’s protocol. Assay was performed at least 12 hr after the ultrasound treatment.

### Live cell assay

Calcein AM (Sigma Aldrich) dye was used to check the cell viability according to manufacturer’s protocol. Cells were incubated with dye every day for 15 mins before imaging.

### Calcium indicator dye-based assay

Cell samples were incubated with the calcium indicator dye (4 mM of Cal-520 AM, AAT Bioquest) for 1 hr. Samples were then replenished with fresh culture medium and allowed to stabilize for 30 min prior to ultrasound treatment.

### Immunocytochemistry and fluorescence microscopy

Samples were fixed with 4% paraformaldehyde (Thermofisher Scientific) solution for 10 min and permeabilized with 0.2% Triton X-100 for 5 min. Normal goat serum (2%) was used as a blocking buffer and samples were treated with serum for 1 hr. Samples were then incubated with primary antibodies of rabbit polyclonal anti-Piezo1 (1:200, Novus Biologicals, catalogue no. NBP1-78446) overnight at 4°C followed by treatment with Alexa Fluor-594 secondary antibodies (Thermofisher Scientific). Hoechst dye (1:1000, Thermofisher Scientific) was used to stain cell nucleus. For *in vitro* study, fluorescence and bright field images were acquired using wide-field Olympus live-EZ microscope equipped with photometrics CoolSNAP K4 camera and W1 live-SR spinning disk microscope equipped with photometrics Prime 95B sCMOS camera. For *in vivo* tumor imaging, wide-field Zeiss stereomicroscope was used.

### Tumor organoid formation

Pancreatic tumor tissue was obtained from patients undergoing endoscopic surgical resection or tissue biopsy at National University Hospital, Singapore. All tissue donation and experiments were reviewed and approved by the Institutional Review Board. Tissue samples were collected in Hank’s Balanced Salt Solution and transferred from the hospital at 4°C. Tissues were minced and plated in Matrigel (Corning). Cell culture medium used is as follows: advanced DMEM/F12, HEPES 10mM, Glutamax 1X, A8301 500nM, hEGF 50ng/mL, mNoggin 100ng/mL, hFGF10 100ng/mL, hGastrin I 0.01uM, N-acetylcysteine 1.25mM, Nicotinamide 10mM, PGE2 1uM, B27 supplement 1X final, R-spondin1, Wnt3A.

### CAM tumor treatment with ultrasound

Fertilized chicken eggs were incubated at 37°C with 70% humidity starting from embryonic day 0 (ED 0). A window was made on eggs on ED 3 to prevent the CAM from sticking onto the shell during development. Under sterile conditions, a small incision was made at the pointed end of the egg using a pointed surgical rod or needle, and 4 to 5 ml of albumin was removed through the hole in order to lower the CAM. Egg shell was then cut to form a window of approximate 1cm diameter, exposing the CAM. This window was sealed with Tegaderm transparent film dressing (3M). Eggs were further incubated to ED7 until they were become ready for tumor grafting. GFP transfected stable MDA-MB-231 and GFP transfected (transiently expressed) transformed HEK293T cells were used grafting purpose. For grafting onto CAM, tumor cells were trypsinized, washed by PBS, and re-suspended in pre-chilled Matrigel (Corning). Next, eggshell was further cut to form a larger window. By gently touching the upper periderm layer of the CAM with an autoclaved glass rod, a slightly bruised area was formed, and one million cells in 50 μl Matrigel were inoculated drop-wise onto bruised area. After inoculation, the window was sealed again with film dressing. On ED10, tumors formed were imaged using stereomicroscope (optical and fluorescence) before treating with the ultrasound. Fertilized eggs containing tumors were subjected to ultrasound for 2 hr. Procedure was done for two consecutive days ED10 and ED11. Tumor imaging was again performed on ED12 to compare the tumor growth. Only tumors with viable embryos were included in data analysis.

### Ultrasound coupling

The primary objective of the sound coupling was to create tensile stress in the membrane that is relatively constant for time periods on the order of one second while only causing acceptable friction at cell-cell adhesions thereby limiting adverse heat production. This achieved with wavelengths much larger than a cell diameter. The constancy criterion requires the acceleration forces to be comparable to the resistance of pressure against the tissue, limiting the excitation to below 200 kHz. The wavelength at 200 kHz in tissue is some 7.5mm, hence friction heating is not a limiting factor.

### Ultrasound transducer

We used ring transducers with diameters of 16mm and 25mm made of PZT4 material that can be effectively driven up to 100s of (Beijing Ultrasonics 25×10×4 and 16×8×4 piezoceramic rings).^34^ They can travel large amplitudes of around 1.5 μm at low frequencies. An aluminium cone is mounted on the rings to widen the field to some 5 cm diameter and attenuate the amplitude, in order to form plane waves. Aluminium has good coupling to the PZT material but poorly matches the water bath due to its very different *acoustic impedance*. We added epoxy and silicon rubberized coats to improve the emission.

In order to avoid standing waves and the near field of the transducer, we use a 2λ deep 37°C water bath between the top of the transducer and the specimen. The specimen itself was floated on the water surface inside a standard multi well plate (Thermo Scientific Nunclon Delta Surface 96 well plates). Those are needed to be tape sealed in order to prevent water creep under ultrasound irradiation. The well plates was suspended on a PET foil with a rectangular opening at the bottom on the plate. Polymer and glass cover slips both showed minimal attenuation of the beam strength. To eliminate standing wave patterns that create spatial intensity fluctuations, the transducer base was suspended and mounted on absorber foam. The influence of surface reflections was minimised by sweeping the frequency 5% to 15%, leading to an about 80% homogeneity in the centre of the sound cone. This finally allowed for controlling the power emitted via the driving voltage of the transducers. We used both a resistor networks and a class A amplifier with interchangeable results. Despite the shielded wiring, radiated radio noise from the transducers remained considerable and the incubator cabinet was used as a shielding capsule.

### Ultrasound generators

All primary signals and sequences were generated by ArduinoDue microcontrollers. This allowed altering duty cycles, sequences, and duration swiftly. The analog setup, used to calibrate the entire system, did output the waveform via the Arduino’s digital-to-analog-converter, then amplified via a medium power amplifier (LM 675 from Texas Instruments) with a symmetric ± 24V supply. This drives a high frequency ferrite ring core (Richco Ferrite Ring Toroid Core, 31.5 × 19.3 × 8mm) isolation 1:5 transformer feeding up to 200V amplitude to the transducers.

For higher power levels, the generator was operated as a switching power supply from a single 24V source. The 3.3V logic of the controller shuttering two transistor switches (Infineon IRFP260MPBF 50A, 200V N-Channel MOSFET) which drove each input of the same transformer. Pull-up was provided by 6, 8, or 10 Ohm power resistors with 50W loss ratings. The transformer inputs were clamped by varistor (EPCOS Varistor 8nF 20A 56V) for the switching design and by diodes (1N5408) to the supply power for the amplifier design. The tunable range was kept within 50% bandwidth of the design frequency of the transducer, i.e. a 30 kHz generator was used from some 22.5kHz to about 37.5kHz. The controller outputs the waveform sample at 1 M sample/second. Exact 30 kHz can be output by storing 3 full waves in 100 samples which then are output 10000 times a second. A soft start and stop is provided by ramping up and down the amplitude over 30 waves. i.e. the same time-base provides accurate on-off cycles.

