## Supplementary file for "Ultrasound-mediated mechanical forces selectively kill tumor cells"

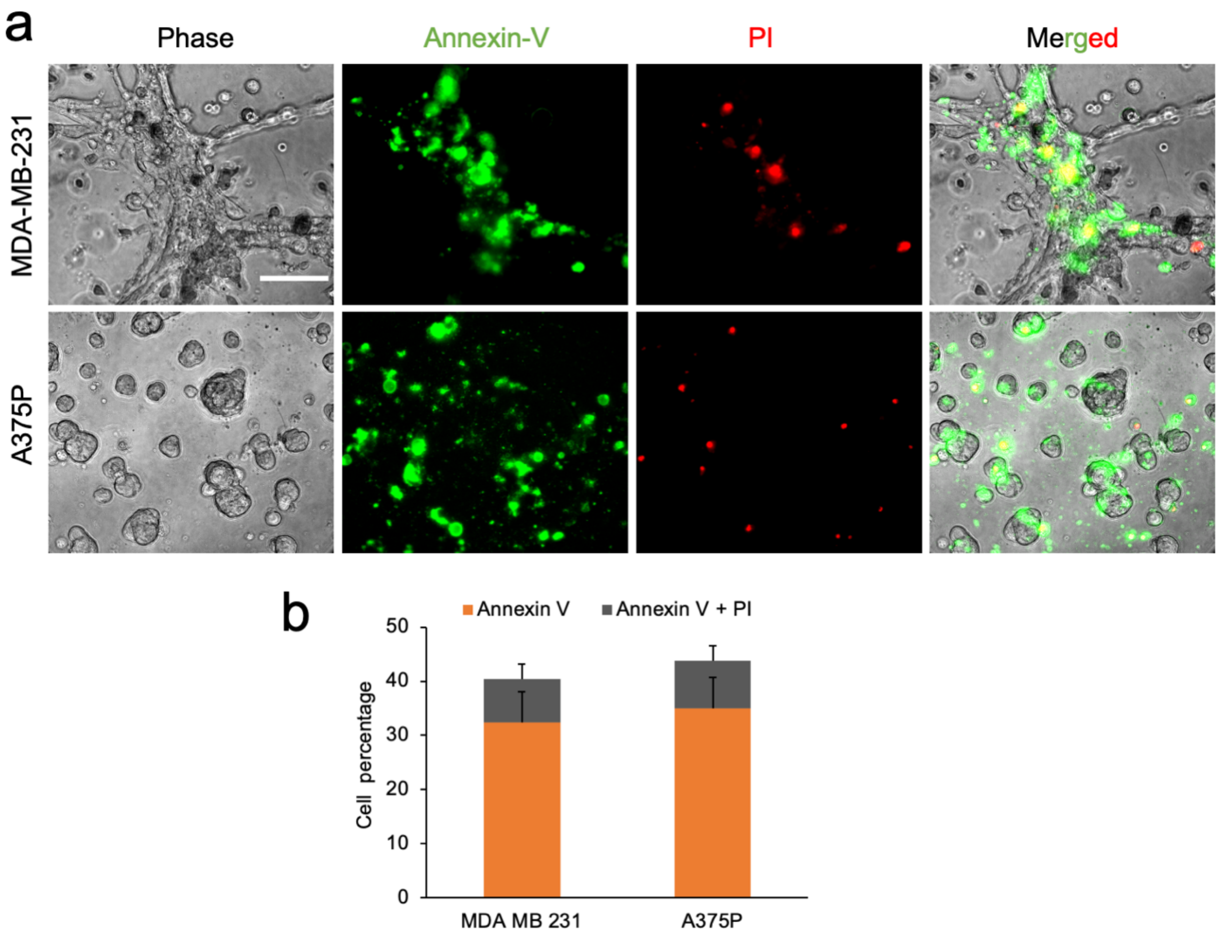


**Figure S1. Ultrasound mainly promotes the apoptosis in tumor cells (a)** Representative images displaying apoptotic (annexin V) and necrotic (propidium iodide) cells after the treatment, scale 100 μm. **(b)** Bar diagram demonstrating percentage of cells showing apoptosis and necrosis with treatment, n>650 cells.

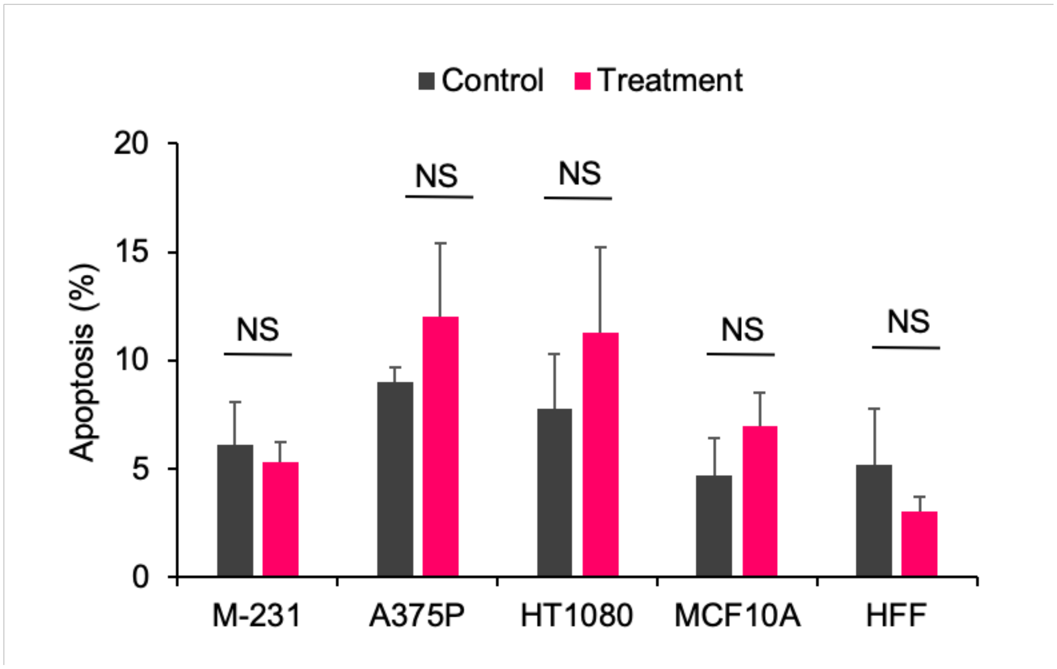


**Figure S2. Ultrasound at 120 kHz frequency do not cause tumor cell apoptosis**. Bar diagram illustrating apoptosis level in tumor (MDA-MB-231, A375p, HT1080) and normal cells (MCF10A, HFF) with and without ultrasound treatment. n>1000 cells, data are representative of two independent experiments, Annova test, $p$ <0.05, no significance.

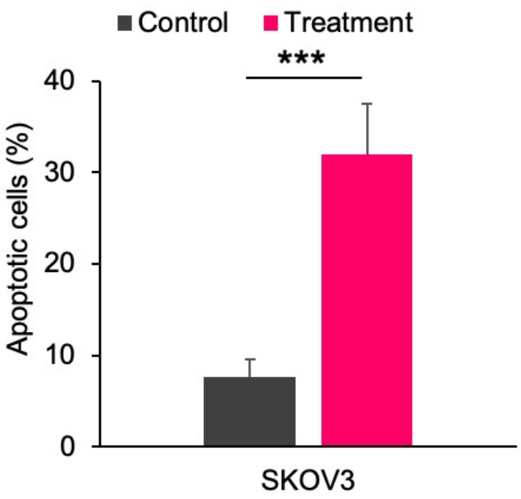


**Figure S3. Ultrasound induces apoptosis in tumor cells from different tissue origin.** Bar diagram showing apoptosis level in SKOV3, ovarian adenocarcinoma with and without ultrasound treatment. n>750 cells, student t-test, $p$***< 0.001

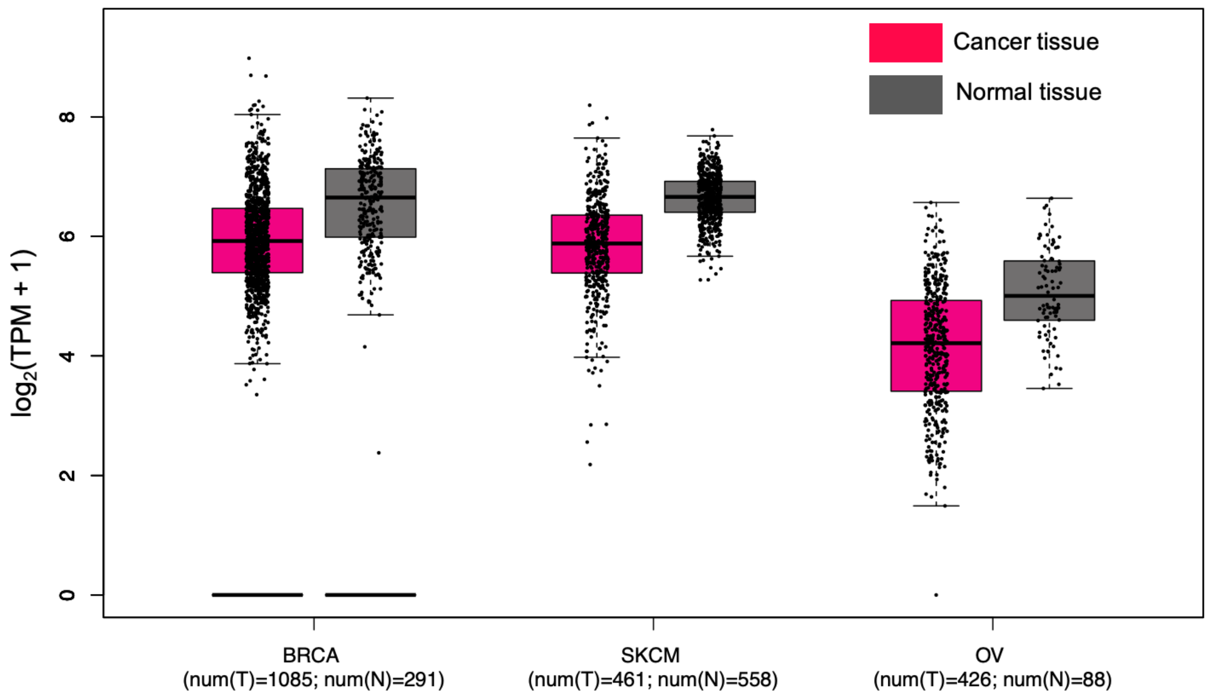


**Figure S4.** TCGA+GTEx data illustrating the expression level of Piezo1 gene in breast carcinoma (BRCA), skin melanoma (SKCM) and ovarian adenocarcinoma (OV) and their respective normal tissues. No significant difference in *p* values. Analyzed by GEPIA2 tool.

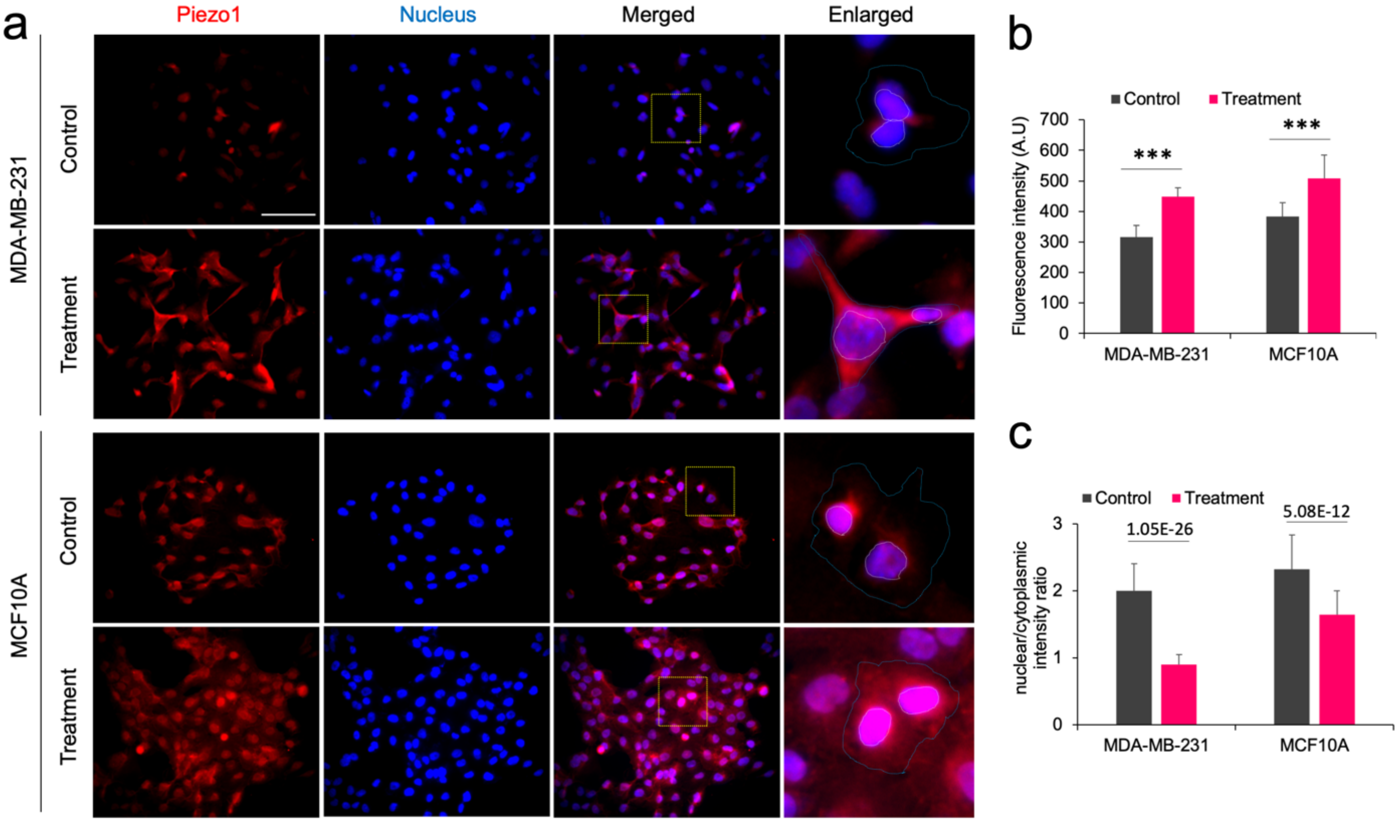


**Figure S5. Ultrasound promotes Piezo1 localization to the plasma membrane in tumor cells.** Representative images displaying Piezo1 expression in MDA-MB-231 and MCF10A cells with and without treatment. Scale bar: 100 µm. **(b)** Bar diagram showing Piezo1 fluorescence intensity profile in tumor and normal cells with and without treatment, n>10 image fields, Annova test, $p$***<0.001. **(c)** Bar diagram illustrating the nuclear/cytoplasmic intensity ratio of Piezo1 in tumor and normal cells with and without treatment, n>55 cells, Annova test, $p$***<0.001.

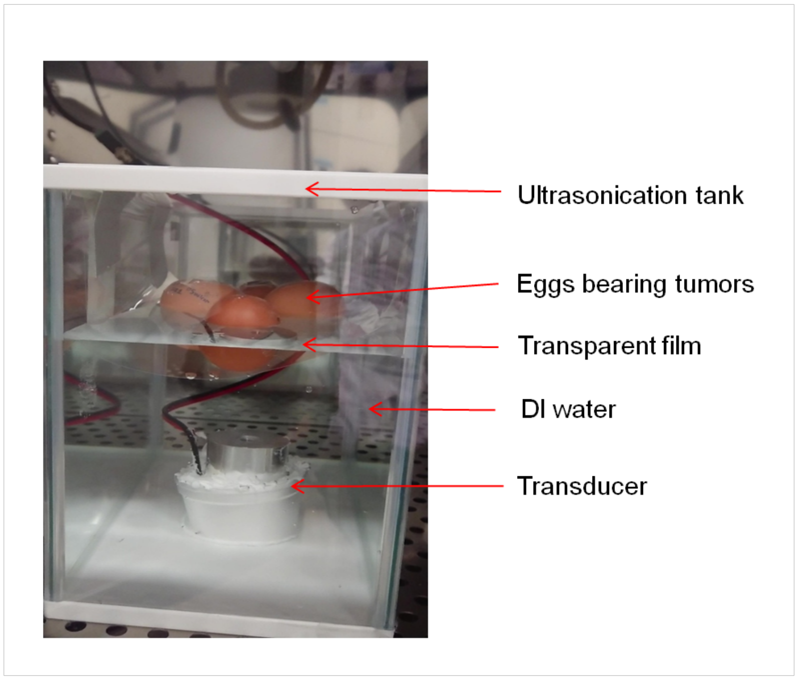


**Figure S6. Picture displaying ultrasound tank containing fertilized eggs**. Fertilized eggs bearing tumors were half submerged in water using transparent film and mounted above the ultrasound transducer for exposure.

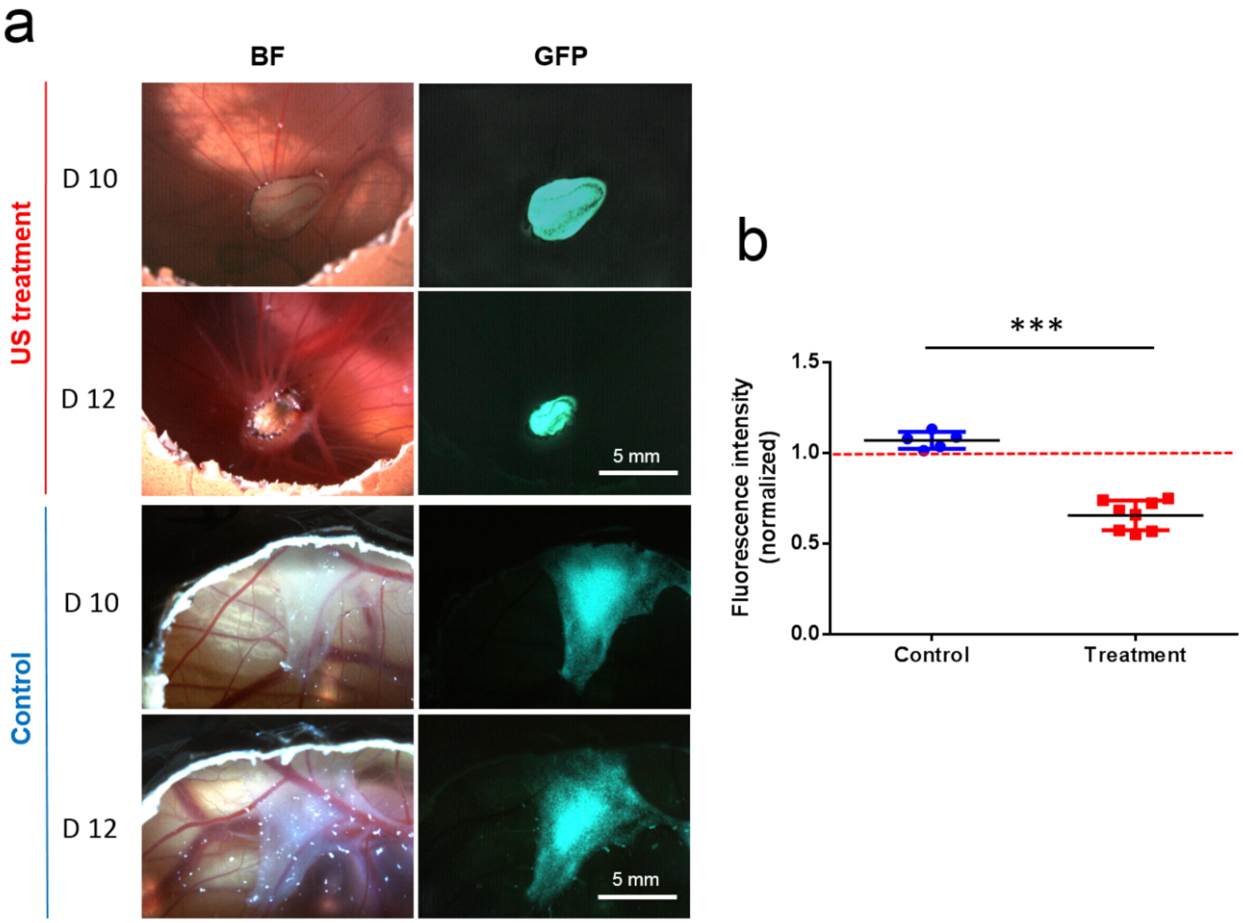


**Figure S7**. **Ultrasound causes reduction in HEK293T tumor growth. a)** Top two panel rows display bright-field and GFP images of tumor before (D10) and after two rounds (D12) of treatment. Bottom panel rows show bright-field and GFP image of tumor without ultrasound treatment on D10 and D12. **b)** Graph showing fluorescence intensity of tumors with and without US treatment on D12. Baseline intensity is shown by red dotted line. (n=5 for control and n= 8 for treatment), data are representative of two independent experiments, $p$***< 0.001, student t-test.

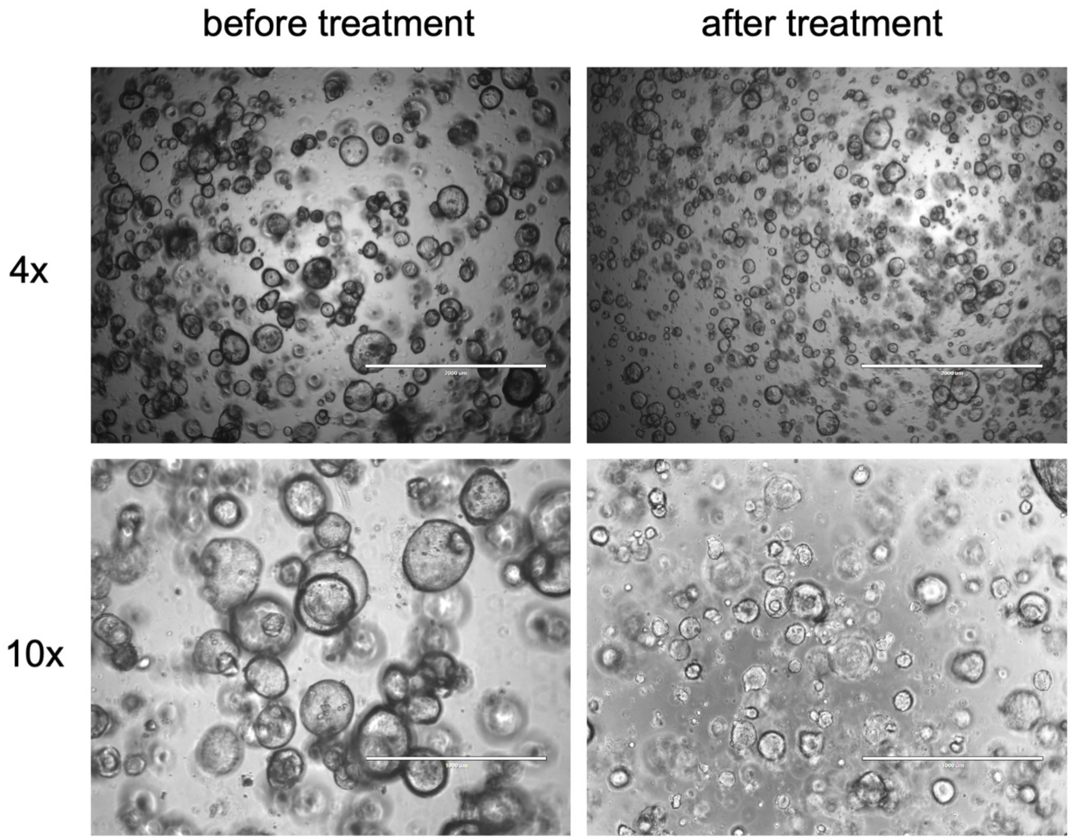


**Figure S8. Ultrasound treatment disrupts pancreatic tumor organoids.** Representative images showing organoid morphology before and after ultrasound treatment at 4x and 10x magnification. Scale bar: 2 mm (4x) and 1 mm (10x)
